# Identification of developmentally important genes in *Silene latifolia* through chemical genetics and transcriptome profiling

**DOI:** 10.1101/2021.01.25.428076

**Authors:** Václav Bačovský, Radim Čegan, Eva Tihlaříková, Vilém Neděla, Vojtěch Hudzieczek, Lubomír Smrža, Vladimír Beneš, Roman Hobza

**Author notes:** **Corresponding author** Roman Hobza.

## Abstract

Dioecious plants possess diverse sex determination systems and unique mechanisms of reproductive organ development; however, little is known about how sex-linked genes shape the expression of regulatory cascades that lead to developmental differences between sexes. In *Silene latifolia*, a dioecious plant with stable dimorphism in floral traits, early experiments suggested that female-regulator genes act on the factors that determine the boundaries of the flower whorls. To identify these regulators, we sequenced the transcriptome of male flowers with fully developed gynoecia induced by rapid demethylation in the parental generation. As the hermaphrodite flower trait is holandric (transmitted only from male to male, inherited on the Y chromosome), we screened for genes that are differentially expressed between male, female, and hermaphrodite flowers. Dozens of candidate genes that are upregulated in hermaphrodite flowers compared to male and female flowers were detected and found to have putative roles in floral organization, affecting the expression of floral MADS-box and other genes. Amongst these genes, eight candidates were found to promote gynoecium formation in female and hermaphrodite flowers, affecting organ size, whorl boundary, and the expression of mainly B class flower genes. To complement our transcriptome analysis, we closely examined the floral organs in their native state using a field emission environmental scanning electron microscope. Our results reveal the principal regulatory pathways involved in sex-specific flower development in the classical model of dioecy, *S. latifolia*.

## Introduction

Sex determination in plant and animal species is driven by environmental cues (e.g., temperature affects sex ratio in turtles) and/or by genetic factors located on the sex chromosomes (Abbott *et al*, 2017). Dominant gene(s) on sex chromosomes control the development of female and male organs and influence specific properties beyond sexual organogenesis, such as size, weight, color, and behavioral functions (Hurst, 1996; Tanurdzic & Banks, 2004). Several studies in plants, namely, papaya (*Carica papaya*) (Ming *et al*, 2007; Iovene *et al*, 2015), persimmon (*Diospyros kaki*) (Akagi *et al*, 2016, 2014), poplar (*Populus trichocarpa*) (Tuskan *et al*, 2006), garden asparagus (*Asparagus officinalis*) (Harkess *et al*, 2020, 2017) and kiwifruit (*Acitinidia chinensis*) (Akagi *et al*, 2019, 2018), have shown that sex determination involves complex molecular pathways. In garden asparagus (Harkess *et al*, 2017) and kiwifruit (Akagi *et al*, 2019), the male activator and female suppressor genes localized on the Y chromosome support the “two-gene” sex determination model that posits two separate genes (usually in one linkage group, male and female sterility factors) in the evolution of dioecy (Cronk & Müller, 2020). In poplar and persimmon, on the contrary, the sex determination is controlled by single locus (Akagi *et al*, 2014; Müller *et al*, 2020), implying that diverse upstream regulators control dioecy in plants, reflecting their polyphyletic origin. The anther regulator gene in the hexaploid Japanese persimmon (*Diospyros kaki*) also maintains an alternative epigenetic state whose specific expression may result in sex reversion (Akagi *et al*, 2016). This study showed that sex determination is also controlled by epigenetic factors, which may differ between the X- and Y-linked alleles, as supported by studies in poplar (Bräutigam *et al*, 2017) and white campion (Bačovský *et al*, 2019; Luis Rodríguez Lorenzo *et al*, 2020; Rodríguez Lorenzo *et al*, 2018). To date, no positive female regulator gene has been described in any dioecious plant and it is recognized that such knowledge would shed light on the origin of the sex-determining chromosome region.

*Silene latifolia* is an excellent plant model for studying sex chromosome evolution, sex determination (Hobza *et al*, 2018), Y chromosome degeneration (Charlesworth, 2016; Vyskot & Hobza, 2015), and dosage compensation (Muyle *et al*, 2017; Wright *et al*, 2016). Yet, identification of the sex-determining genes in this dioecious plant is hampered by the locus complexity of the non-recombining region, paired with the abundance of transposons (Hobza *et al*, 2018, 2015). Early studies examined epigenetic regulation of gynoecium development in *S. latifolia* by treatment with 5-azacytidine (5-azaC), which incorporates into DNA and inhibits DNA methyltransferases, causing DNA hypomethylation in a reversed manner. The authors proposed three possible mechanisms for gynoecium development and suggested that epigenetically modified genomic regions undergo holandric inheritance (Janoušek *et al*, 1996, 1998). Several studies used biological treatment and smut fungus infection to cause an effect similar to that of 5-azaC treatment (Janoušek *et al*, 1996), promoting female flower masculinization and male feminization (Kazama *et al*, 2005; Zemp *et al*, 2015; Kawamoto *et al*, 2019). Although such experiments proved to be efficient for studying the developmentally important functions of flower-related genes, the atypical developmental changes were not heritable. Thus, the transcriptome expression profiles were not altered in the long term to reveal a complete set of candidate alleles promoting flower organization (Kazama *et al*, 2005; Hobza *et al*, 2018). Recently, based on the analysis of asexual flowers, it was suggested that smut fungus infection will have the same effect on flower development as the genes located in the stamen promoting factor (SPF) region (Kawamoto *et al*, 2019). Yet, none of these genes have been identified, and further evidence is needed to understand the nature of sex determination and dosage compensation in this model plant (Wright *et al*, 2016; Ponnikas *et al*, 2018; Muyle *et al*, 2017) and to identify female- and male-specific genes.

Treatment with epigenetically active chemical agents (chemical genetics) alone and in combination is a common practice in transgenerational studies examining epigenetic regulation of various traits (Zhang *et al*, 2013; Cruz & Becker, 2020). These traits are established through a new variety of differentially methylated regions (Zhang *et al*, 2018; Bartels *et al*, 2018), leading to allele expression repatterning and new phenotypic characteristics transmitted to the next generation (Baubec *et al*, 2014; Reinders *et al*, 2009; Pecinka & Liu, 2014). Such treatment is often used if mutants are not available or if specific interference with DNA in order to obtain new phenotypes is required (Pecinka & Liu, 2014). Like mutagenesis, chemical genetic approaches may result in pleiotropic effects and phenotypes that are not identical to genetic mutations (Baubec *et al*, 2010; Pecinka & Liu, 2014). To examine the pleiotropic effects caused by chemical genetics, next-generation high-throughput RNA sequencing (RNA-seq) and transcriptome profiling are fundamental tools to understand the functional genomic elements in various species (Wang *et al*, 2009; Shendure *et al*, 2017). In many plants, RNA-seq analysis has led to characterization of multiple regulatory pathways of important traits such as flowering (Glazinska *et al*, 2017; Quan *et al*, 2019; Hu *et al*, 2020), hybrid vigor (Howlader *et al*, 2020; Li *et al*, 2009), and resistance to biotic or abiotic stress conditions (Kumar & Trivedi, 2016; Kang *et al*, 2020). Thus, transcriptome profiling provided valuable information about the expression of important genes in many species (Wang *et al*, 2009; Mortazavi *et al*, 2008; Kumar & Trivedi, 2016; Shulse *et al*, 2019).

In this work, we aimed to address two important questions concerning sex determination in *S. latifolia* using a combination of RNA-seq transcriptome profiling and precise phenotyping: First, what regulatory pathways are involved in gynoecium development in *S. latifolia*, and second, how are the principal regulatory pathways involved in flower development linked to sex-specific gene expression? Using precise phenotyping with a field emission environmental scanning electron microscope (FE-ESEM), we describe how the atypical development of the gynoecium in males affects hermaphrodite fertility and placenta organization. To identify the gene regulatory pathways involved in gynoecium development, we gathered the transcriptomes of males, females, and hermaphrodites and compared their expression profiles. Using this approach, we identified previously unknown important regulatory sex-linked and autosomal genes involved in female (gynoecium) development and transcription factors involved in flower organization. We discuss the possible function of newly detected sex-linked genes in the context of previous results and hypothesize that they have a role in the classical ABC(DE) model of flower development in *S. latifolia.* The transcription profiles show stable inheritance once impaired, confirming previously hypothesized suppression of developmentally important genes in *S. latifolia* flower development in male flowers. We show how the upstream regulatory pathways affect downstream and MADS-box gene expression in flower development.

## Methods

### Treatment data, drug concentration and RNA extraction

Seedlings of *S*. *latifolia* inbred population U15 (seeds owned by Institute of Biophysics of the Czech Academy of Sciences), made by 15 generations of full-sib mating, were used as a parental population for all drug experiments (SI Appendix, Supplemental Methods). Efficient drug concentration was estimated based on the literature (Baubec *et al*, 2009; Lechner *et al*, 1996) and an *in vitro* drug treatment assay (SI Appendix, Figs. S1a, S2, S3). Mixture injections were repeated every 3 days for 2 weeks after the first occurrence of flowering (SI Appendix, Fig. S1b). Seeds derived from self-pollinated hermaphrodite flowers and full-sib mating crosses in F1 and F2 were used to test the inheritance pattern of the hermaphrodite phenotype. RNA was isolated in biological triplicates from flowers of stage 11 based on Zluvoval *et. al*. (52; SI Appendix, Supplemental Methods).

### Flower phenotyping

Flowers were opened with dissecting needles, and petals and sepals were carefully removed for final imaging and ovule and placenta examination (SI Appendix, Supplemental Methods). A pollen staining protocol (Peterson *et al*, 2010) was followed with minor modification as described (SI Appendix, Supplemental Methods). For morphological characterization of flowers in their fresh state free of any microstructural changes or artifacts, a field emission environmental scanning electron microscope equipped for advanced control of thermodynamic conditions inside the specimen chamber (Neděla *et al*, 2020), the wide field aperture detector (WFAD) for ESEM (newly developed for the purpose of this research at the ISI ASCR), and the ionization secondary electron detector with an electrostatic separator (ISEDS) were used (Neděla *et al*, 2018). For FE-ESEM, sepals and petals were first cut off from stage 11 flowers in the same way as for phenotyping and RNA isolation (SI Appendix, Supplemental Methods).

### RNA-seq library construction and sequencing

Transcriptome sequencing libraries were prepared from total RNA with the Illumina TruSeq mRNA Library Prep Kit at the Genomics Core Facility (Genecore, EMBL, Heidelberg). The sequencing of the libraries was conducted at the Genecore on Illumina NextSeq500 instrument with 75-bp paired-end reads.

### RNA-seq data analysis

The quality of sequencing reads was inspected in FastQC (http://www.bioinformatics.babraham.ac.uk/projects/fastq). The *S. latifolia* U10 reference transcriptome (Zemp *et al*, 2015) was verified by the Benchmarking Universal Single-Copy Ortholog (BUSCO) pipeline (Simão *et al*, 2015). Read trimming on quality (Q30) and sequencing adaptor removal were done with Trimmomatic −0.32 (Bolger *et al*, 2014). Cleaned reads from each library were pseudoaligned to the *S. latifolia* reference transcriptome (Muyle *et al*, 2012), which originated from the U population, using Kallisto (version 0.45.0) (Bray *et al*, 2016) with default parameters and number of bootstrap samples set to 100. The Kallisto index for reference fasta files was created with a k-mer length of 19. Transcript abundance estimations from Kallisto were used as input for differential expression analysis using the Bioconductor DESeq2 package (version 1.24.0) (Love *et al*, 2014). The generalized linear model approach was applied to all comparisons. Transcripts were considered as differentially expressed when the adjusted *P* value was <0.001 and log2 fold change was >±1. Transcripts were annotated through comparisons with the Swissprot and UniprotKB complete proteome databases using the Trinotate pipeline (Table S1) (Bryant *et al*, 2017). Gene Ontology (GO) enrichment among selected transcripts was done using Trinotate and GOseq bioconductor package (Young *et al*, 2010). Clustering of differentially expressed genes (DEGs) was done by the bioconductor coseq package (SI, Supplemental Methods). All R scripts were run in R studio, and all software and tools used are listed in supplementary notes (SI Appendix, Supplemental Methods). Repetitive sequences were identified in the reference transcriptome by blast searches (with e-value 1E–20) against the *S. latifolia* repeat database (Macas *et al*, 2011).

### Identification of sex-linked transcripts

Putative sex-linked transcripts were identified in the *S. latifolia* reference transcriptome (Zemp *et al*, 2016) by blast-n searches against previously published sets of sex-linked transcripts (Zemp *et al*, 2016; Papadopulos *et al*, 2015; Chibalina & Filatov, 2011; Bergero & Charlesworth, 2011; Muyle *et al*, 2012). For blast searches, blast-n with e-value 1E–40 from blast-all 2.2.26 was used (ftp://ftp.ncbi.nlm.nih.gov/blast/executables/). As sex-linked transcripts we considered those with at least one hit against mentioned datasets (Table S2).

## Results

### Epigenetic chemical treatment turns males into hermaphrodites

Similarly to previous experiments (Janoušek *et al*, 1996, 1998), we induced hermaphrodite flower development in male individuals of *S. latifolia* using two functionally diverse groups of chemical drugs (Nowicka *et al*, 2019): i) zebularine (ZEB), a non-methylable cytidine analog, and ii) sodium butyrate (SB), a histone deacetylase inhibitor. First, both drugs were screened for their optimal dose concentrations (SI Appendix, Supplemental Note S1, Fig. S1a). Second, the optimal dose concentrations were injected into adult *S. latifolia* plants before flowering, and finally, the hermaphrodite flower trait was followed for two generations (SI Appendix, Figs. S1b, S2, S3).

We examined the effect of optimal doses on adult plants for ZEB and SB separately and in combination (40 μM ZEB, 2.5 mM SB; Fig. S3a–c; Supplemental Note S2). As expected from the *in vitro* assay (SI Appendix, Supplemental Note S1), ZEB and SB separately turned 20–30% of males in the population (*n*=128) into hermaphrodites (SI Appendix, Fig. S3d). The combination of both drugs induced hermaphrodite flowers in almost 40% of males (SI Appendix, Fig. S3d). The control group of males and females in the *in vitro* assay contained no individuals with prolongated carpels or prolongated stamens. Next, we used flowers with both carpels (gynoecium) and stamens as pollen donors in the crossing experiment (SI Appendix, Fig. S2a). Again, almost 40% of the males in the F1 population were hermaphrodites (termed HF1), and all males in the F2 population (termed HF2) were hermaphrodites (SI Appendix, Fig. S3e). As a control, we used non-treated (NT) males as pollen donors and pollinated treated females in the F0, F1, and F2 generations (directly after treatment or in the HF1 and HF2 generations). No hermaphrodites resulted from such crosses (SI Appendix, Fig. S2b); hence, the hermaphrodite flower trait was transmitted only through the male germline (SI Appendix, Fig. S2a). We observed four different stages of hermaphrodite flowers (HF_I-IV_) and types of carpel (SI Appendix, Fig. S2c). HFI contained strongly reduced ovary and atypically prolongated stigma. HF_II_ consisted already of one developed stigma as well as suppressed stigma and ovary without placenta. HF_III_ and HF_IV_ possessed one to four stigma, but the size of placenta was decreased compared to NT females (SI Appendix, Fig. S2c). The inheritance of these flower traits was increased when more perfect flower types (functional ovary and stigma) of hermaphrodite stage HF_III-IV_ were used as pollen donors (SI Appendix, Fig. S3d, e).

### Hermaphrodite flowers contain defective carpels and fewer ovules

We studied detailed phenotypic changes caused by ZEB and SB using the combination of large-field and ISEDS morphology-sensitive detectors of secondary electrons (Neděla *et al*, 2020, 2018). We examined male and female flowers in the hermaphrodite progeny using FE-ESEM and found differences at the cellular level in flower organization, placenta, and pollen viability (Fig. 1a–h, SI Appendix, Fig. S4a–h). NT male flowers contained functional stamens and minor suppressed gynoecia (Fig. 1a–c). The hermaphrodite flower type HF_III_ possessed, apart from stamens, a fully fertile gynoecium (Fig. 1d), consisting of a suppressed stigma (Fig. 1e), ovary, and stigma (Fig. 1f), having phenotypically functional anthers (Fig. 1g) similar to the NT male (Fig. 1c). Hermaphrodite flowers possessed significantly higher numbers of aborted pollen compared to NT (*P* < 0.001) (SI Appendix, Fig. S5a–i, Supplemental Note S3). A subtype of hermaphrodite flower termed HF_IIIa_ had phenotypically normal stamens and an ovary containing only one stigma and suppressed stigma (Fig. 1h). All hermaphrodite flowers also significantly differed in the number of ovules (*P* < 0.001) and, in some cases, the placental organization (HFIIIa flowers had marginal placentas, whereas more perfect hermaphrodite flowers possessed free central placentas similar to NT females) (SI Appendix, Fig. S6a-d, Fig. S7a-e, Supplemental Note S3). Female carpels in the HF1 and HF2 generations were similar to NT (SI Appendix, Fig. S4a-h). The flowers consisted of functional ovary (SI Appendix, Fig. S4a, c, e) with 5-6 stigmas (SI Appendix, Fig. S4a, b, e, h), suppressed stamens in NT females (SI Appendix, Fig. S4d), and atypical prolongated stamens in HF1 and HF2 females (SI Appendix, Fig. S4g).

**Fig. 1.**
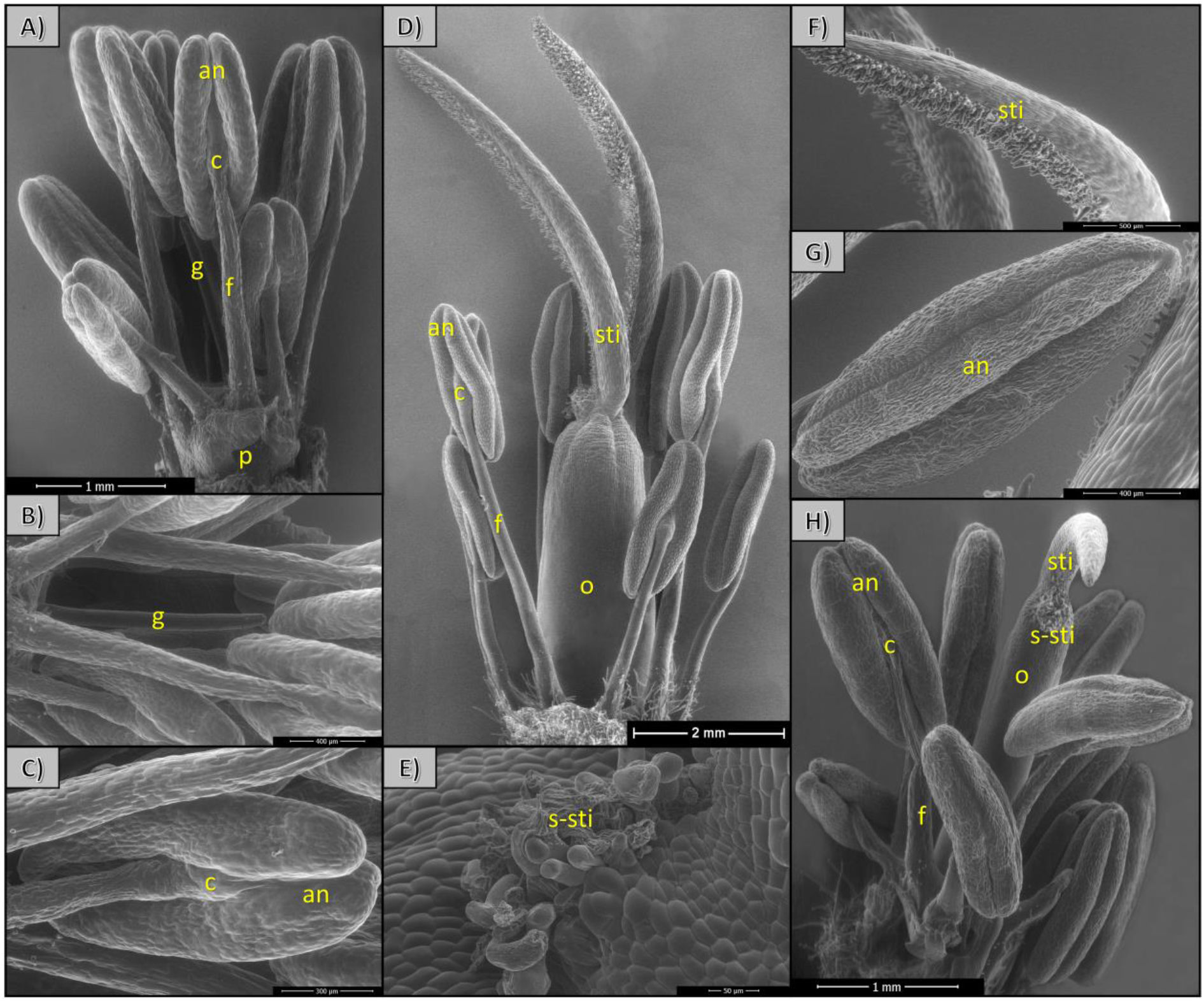
Detailed phenotyping of gynoecium organs in male and hermaphroditic flowers. (**A**) Stage 11 WT male flower, with (**B**) a suppressed gynoecium and (**C**) fully fertile anther. (**D**) Stage 11 fertile hermaphroditic flower type III, with (**E**) a suppressed stigma, (**F**) fully developed stigma and (**G**) anther. (**H**) Stage 11 fertile hermaphroditic flower subtype IIIa. Note the differences between the suppressed gynoecium in males (**B**) and fertile gynoecium consisting of an ovary and stigma in the hermaphroditic flower (**D**, **H**). Anther (an), anther connective (c), filament (f), pedicel (p), gynoecium (g), stigma (sti), ovary (o), suppressed stigma (s-sti).

### Differential gene expression analysis in hermaphrodites and RNA-seq data validation

We generated RNA-seq data to quantify changes in gene expression patterns between *S. latifolia* males, females, HF1, and HF2 (Fig. 1a,d; SI Appendix, Fig. S4a). For 12 samples (triplicate samples for each variant), we obtained nearly 605 million paired-end raw sequencing reads (Table S3). We observed that 1,283 transcripts from 1,440 had complete BUSCOs (Table S4), and nearly 59% of the searched sequences were found in the Swissprot database. In the reference transcriptome we identified 2,469 potentially sex-linked transcripts using the blast search algorithm (compared to available *S. latifolia* sex-linked gene datasets; Table S2); 72% to 79% of the RNA-seq data were successfully pseudoaligned to the reference transcriptome (Table S3). Genetic distance was examined by principal component analysis, and validated by Cook’s distances and by sample-to-sample distances, differentiating all datasets to three clusters: males, females, and hermaphrodites (and subclusters HF1 and HF2) (SI Appendix, Fig. S8a-c; Supplemental Note S4).

Differential gene expression analysis in pairwise male:female (MvsF), male:hermaphrodite F1 (MvsHF1), male:hermaphrodite F2 (MvsHF2), female:hermaphrodite F1 (FvsHF1), female:hermaphrodite F2 (FvsHF2), and hermaphrodite F1:hermaphrodite F2 (HF1vsHF2) comparisons identified 15,325 DEGs. Among them, we examined 3,782 DEGs in MvsF, 1,809 in MvsHF1, 2,621 in MvsHF2, 3,119 in FvsHF1, 3,465 in FvsHF2, and 529 in HF1vsHF2 (SI Appendix, Fig. S9a-f, Table S5, S6). From these comparisons, 4,392 genes were used for identification of candidate genes involved in flower sex development in GO enrichment analysis and transcriptome profiling.

### DNA and histone methylation-related gene expression stability between sexes

We performed expression profiling of genes (Table S7) related to histone modification (methylation, acetylation), DNA methylation, and DNA damage and stress response, as well as genes regulated directly by DNA methylation (epialleles). Using this profiling, we assessed differences in expression in hermaphrodites compared to males and females (Fig. 2). Comparing the male, female, and hermaphrodite flowers (MvsF, MvsHF1, MvsHF2, FvsHF1, FvsHF2, and H1vsHF2), we found that only several genes had distinct expression patterns between sexes (MvsF) and between floral phenotypes (MvsH1, MvsH2). In these comparisons, the most distinct expression patterns were found for homologs of *AGO4b* (Fig. 2a), *DDM1b*, *SUVH4b*, and *SUVR4* (Fig. 2b); *JMJ13* and *JMJ26* (Fig. 2c); *SOG1* (Fig. 2d); *HAC1c*, *HAF1*, *HDA8*, *HDA9*, *SRT1*, and *SRT2* (Fig. 2e); and *ABI3* and *ERECTA* (Fig. 2f). The largest differences in expression were in DNA damage response and stress-related genes between sexes (MvsF) (Fig. 2d). Other non-differential expression profiles for DNA methylation, histone methylation, histone acetylation, and deacetylation genes showed that expression was stable (Fig. 2a-c). Only 11 genes (DDM1b, SUVH4b, SUVR4, JMJ13, JMJ26, HAC1c, HAF1, HDA8, HDA9, SRT1, and SRT2) may represent regulators that specifically affect mRNA levels of several developmentally important key genes and were downregulated in males compared to hermaphrodites at the studied developmental stage.

**Fig. 2.**
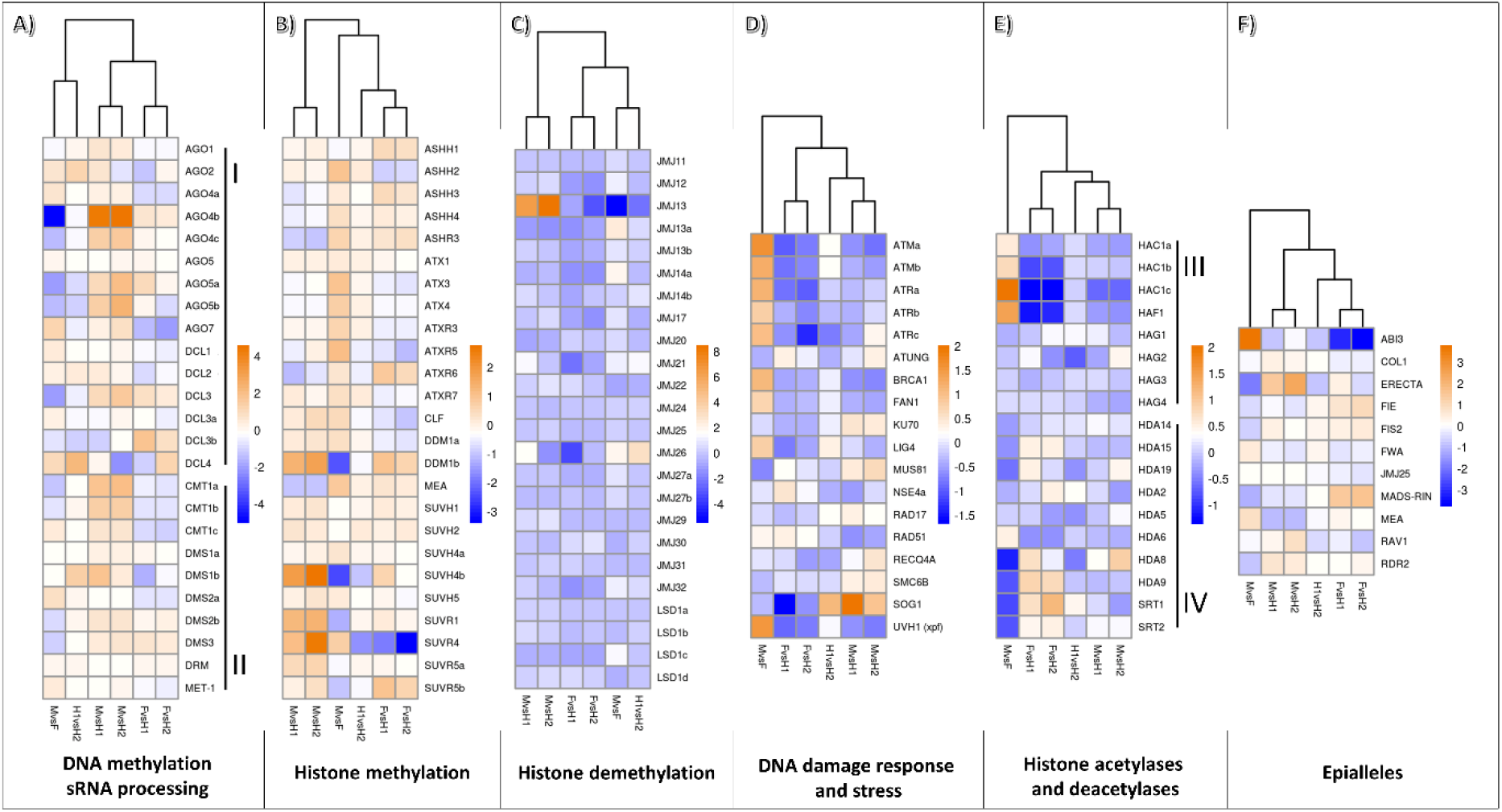
Gene expression patterns related to hermaphrodite development and transcriptome changes after the ZEB and SB treatment. (**A**) Genes involved in DNA methylation (I) and sRNA processing (II), (**B**) genes associated with histone lysine methylation, (**B**) genes associated with histone lysine demethylation, (**D**) genes involved in DNA damage response and stress, (**E**) histone lysine acetylases (III) and deacetylases (IV), (**F**) epialleles or genes regulated by DNA methylation. For corresponding gene IDs and literature sources, see Table S2. Clusters show expression similarity between comparisons.

### Expression of transposable elements *Retand*, *Athila*, and *Copia* is upregulated in hermaphrodites

By blast searches, we identified 158 transcripts from 24 repeat clusters (Table S8). From these clusters, *Copia*, *Athila*, *Peabody*, *Ogre*, *Retand*, *Cacta*, *Sil*, and *LINE* elements were differentially expressed such that expression levels were higher in males compared to females (except for some clusters of *Copia* elements). Compared to males, hermaphrodites (HF1 and HF2) differed mainly in *Retand*, *Athila* and *Copia*, which were upregulated, and *LINE* elements, which were downregulated in HF2 (SI Appendix, Fig. S10, Table S8). These results show that some transposable elements (TEs), namely, *Retand*, *Athila*, and *Copia* families, were deregulated in hermaphrodites and this deregulation was inherited by two generations.

### DEG comparison and expression profiling between sexes yield putative candidates involved in gynoecium development

All differentially expressed transcripts and gene products were classified using GO enrichment analysis into groups involved in biological processes (BP), cellular components (CC), or molecular functions (MF) (SI Appendix, Supplemental Note S5, Figs. S11 and S12). Overall, 362 (4.49%) and 2,296 (2.75%) of GO terms in BP, 83 (3.37%) and 525 (2.34%) in CC, and 372 (2.44%) and 479 (4.28%) of GO terms in MF were up- and downregulated, respectively, in MvsHF1HF2. Among the genes with no classification or known function, 4 (9.9%) and 231 (3.18%) were up- and downregulated, respectively, in MvsHF1HF2 (SI Appendix, Fig. S11, Fig. S12, Table S9). Among the up- and downregulated transcripts in BP, we selected 213 genes that function in floral development (Table S10). These include autosomal and sex-linked genes that were differentially expressed in hermaphrodites and deregulated by the ZEB and SB treatment (Table S10).

We looked for DEGs that were downregulated in MvsF or downregulated in MvsHF1 and MvsHF2 and sex-linked in order to enrich the cluster of gynoecium-promoting candidates. We identified 10 genes that are sex-linked and were differentially expressed in hermaphrodites and 35 genes that are sex-linked and were differentially expressed in both hermaphrodites and females (Fig. 3a). These 45 sex-linked (Table S11) and 333 autosomal DEGs that were downregulated in MvsHF1 and MvsHF2 (Table S12) were searched in Uni-Prot databases, further designated in this study as Group 1. The Uni-Prot searches provided the function for each gene and orthologous sequence in *Arabidopsis*, suggesting similar function in *S*. *latifolia*.

**Fig. 3.**
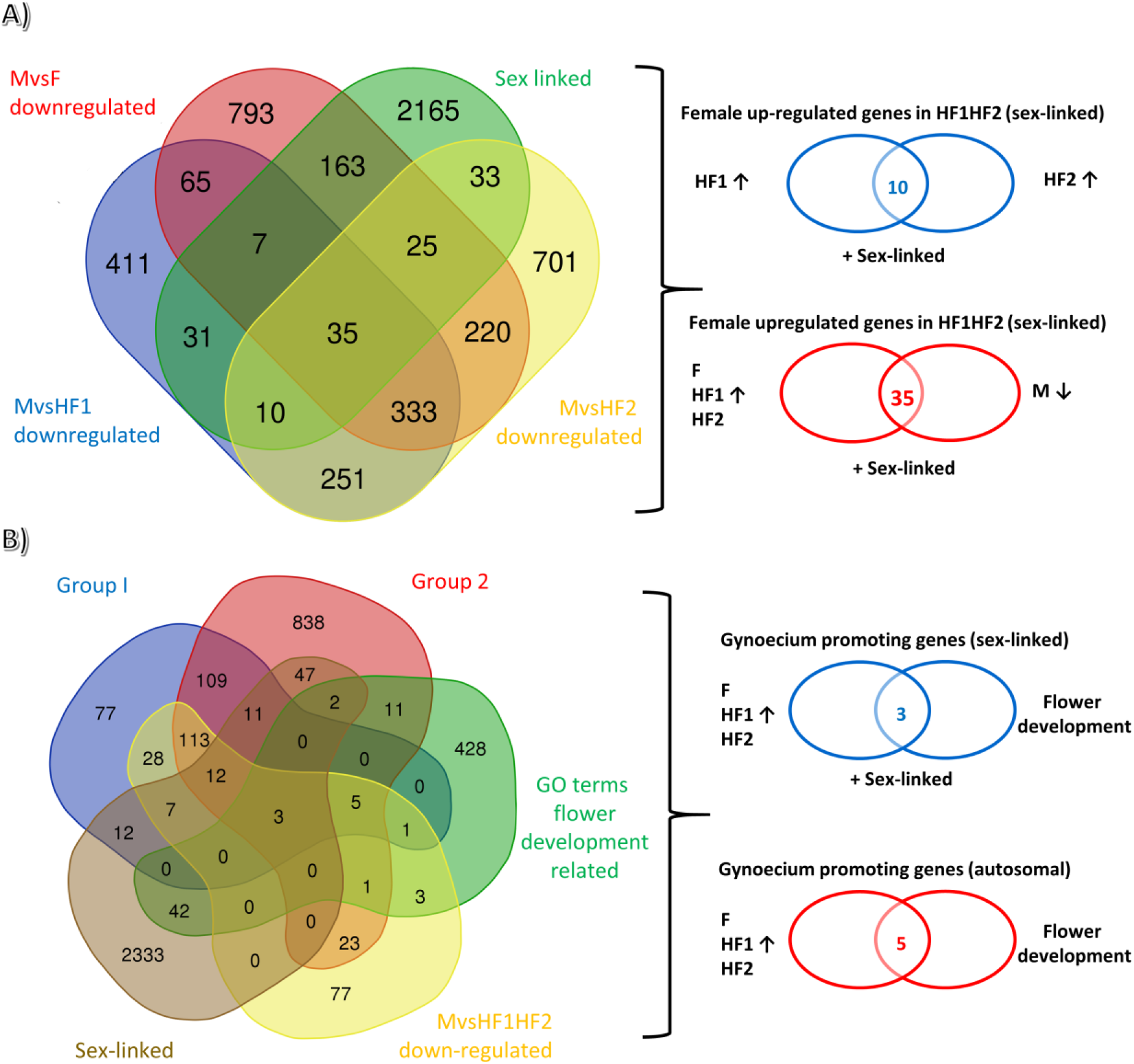
Venn diagrams identifying candidate genes for gynoecium development. (**A**) Genes related to gynoecium development downregulated in males compared to females and hermaphrodites. The gene comparisons are as follows: MvsF downregulated, MvsHF1 downregulated, MvsHF2 downregulated and sex-linked. (**B**) Autosomal and sex-linked candidate genes involved in flower development and gynoecium promotion. The gene comparisons are as follows: Group I (MvsFHF1HF2 downregulated), Group 2 (Cl4; SI Appendix, Fig. S11), GO terms associated with the flower development (Table S7), MvsHF1HF2 downregulated and sex-linked candidates. Non-sex-linked genes enriched in GO terms associated with the flower development pathway were depicted as autosomal.

Clustering analysis identified nine clusters (Cl1-9) distributed into five groups of genes based on their expression: upregulated in hermaphrodites compared to males and females (Cl1-3), having similar levels of expression in hermaphrodites and females (Cl4), strongly expressed in males and having low levels of expression in hermaphrodites and females (Cl5, 6), downregulated in hermaphrodites compared to males and females (Cl7, 8), and having similar levels of expression among sexes (Cl9, SI Appendix, Fig. S13, Table S13). Cl4 was further used to compare developmentally important genes, differentially expressed between males and females, designated in this study as Group2.

### Candidate genes for gynoecium development include essential regulator transcription factors

Genes in Groups 1 and 2 were compared to genes that are sex-linked and were identified based on GO analysis (Table S10) and GO flower development enrichment (Fig. 3b). This comparison newly identified three sex-linked and five autosomal potential candidates, upregulated in hermaphrodites or having similar levels of expression in hermaphrodites and females, for regulating gynoecium development (Fig. 3b, Table S14). The three candidates *GATA18 transcription factor*, known also as *HANABA TARANU* (*HAN or GATA18*); *AP2-like-responsive transcription factor AINTEGUMENTA* (*ANT*); and *Gibberellin 20 oxidase 1* (*GAOX1*; Table S14) were upregulated in hermaphrodites compared to males, with higher levels of expression for *HAN* and *ANT* in females and *GAOX1* in hermaphrodites (Fig. 4a). The five autosomal genes *SQUAMOSA promoter-binding protein-like transcription factor* (*SPL4*), *FRIGIDA-like protein 4a* (*FRL4a*), *Zinc finger protein WIP2* (Z*WIP2*), *Axial regulator YABBY1* (*YAB1*), and *homeobox protein ATH1* (*ATH1*) (Table S14) were strongly expressed in females and hermaphrodites (Fig. 4a). Next, we were interested in the downstream regulatory gene pathways or the expression levels of the binding partners of these proteins (Fig. 4b; SI Appendix, Fig. S14). Within the regulatory pathway were found specifically upregulated genes with putative roles in gynoecium promotion and regulation of MADS genes (SI Appendix, Fig. S14; Table S14).

**Fig. 4.**
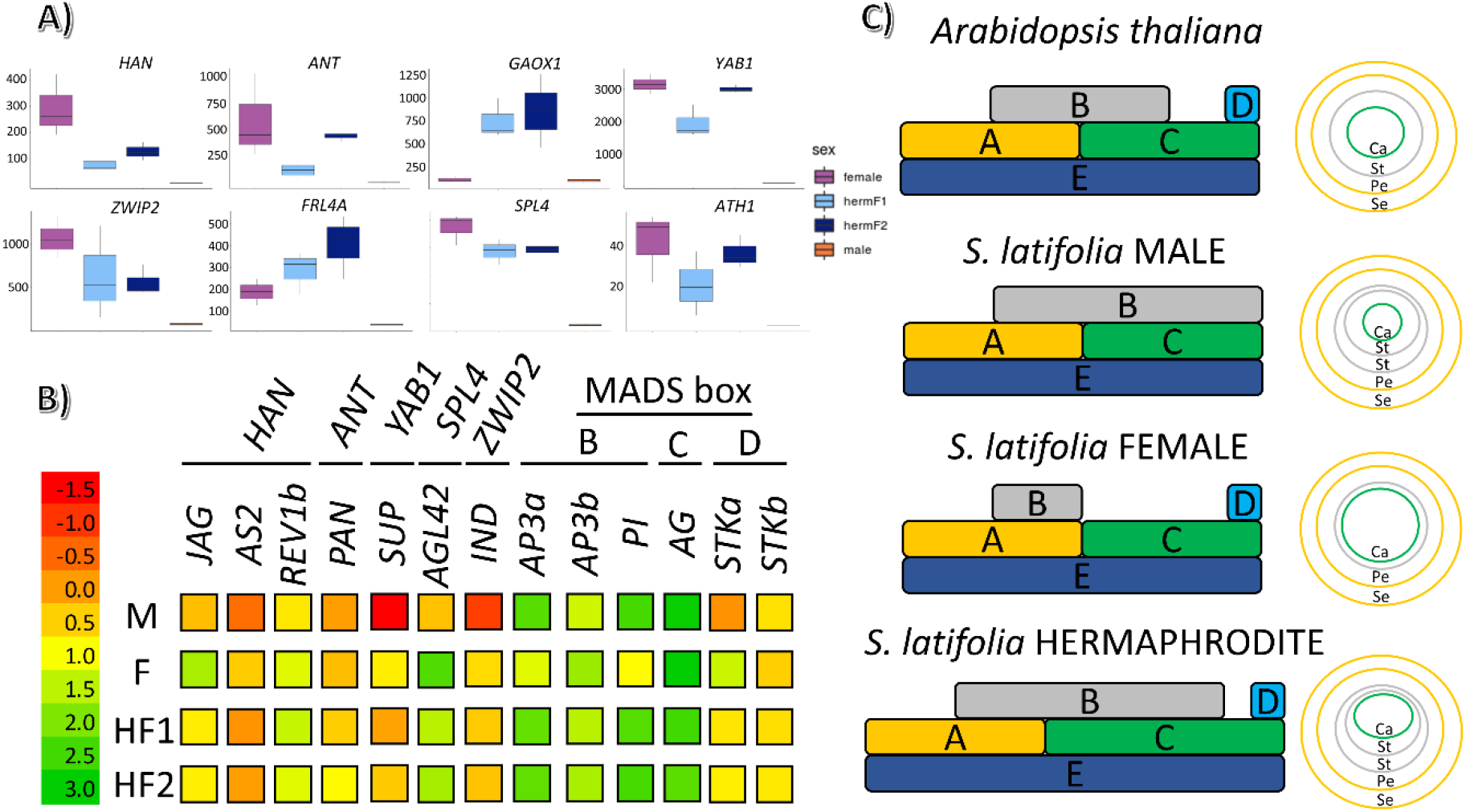
The expression of candidate genes and their regulatory function in relation to the ABC flower development model. (**A**) The relative expression of 8 candidate genes identified in Venn diagram using gene comparison (Fig. 3b). (**B**) The relative expression of high-level regulator and MADS-box B, C, and D genes and their expression changes between sexes. (**C**) ABC(DE) *Arabidopsis thaliana and S. latifolia* flower development model. The model is based on the morphological changes and MADS-box expression changes in hermaphrodite flowers compared to male and female flowers. Whereas wild-type male flowers consist of two whorls for stamen development (10 stamens) and a small reduced carpel, wild-type females possess only one central carpel with 5–6 stigma and 5 rudimentary stamens. The hermaphrodite flower has an enlarged carpel with a reduced number of ovules, 2–3 stigma (usually), and 10 fertile stamens.

According to the classical model of ABC flowering development in *Arabidopsis thaliana*, phenotype and expression analysis in this study suggests that hermaphrodites (genetically males) possessed a larger area in the central whorl (Fig. 4c), keeping the morphology of NT male flowers with strongly enlarged gynoecium (carpel and stigma). This is supported by the fact that males contained 10 anthers and suppressed gynoecium, whereas females possessed only central carpels with five stigmas and five suppressed anthers (Fig. 1, SI Appendix, Fig. S4). Additionally, hermaphrodites had higher expression of class D MADS-box genes, *PAN*, *REV1*, *AGL42*, and *JAGGED*, and eight putative candidates revealed in this study (SI Appendix, Fig. S14). These results show strongly reversed phenotypes from males to androhermaphrodites, which had similar levels of expression for the genes involved in gynoecium promotion (Fig. 4a-c). Additionally, the level of expression of MADS-box genes did not change dramatically as in the case of the upstream regulatory transcription factors (SI Appendix, Fig. S14).

## Discussion

In this study, we show that chemical genetics is a robust tool to analyze and examine novel regulatory pathways in floral development without genomic interference. We influenced the expression of female-specific genes, originally downregulated in males, producing gene reactivation that led to gynoecium promotion in male flowers, turning males into androhermaphrodites. Hermaphrodite flower development was inherited only through the male germline (Fig. S2), confirming previous results (Janoušek *et al*, 1996, 1998). Based on 5-azaC treatment, it was suggested that female flower suppression in *S. latifolia* males is regulated by methylation of specific sequences (Janoušek *et al*, 1996; Bernasconi *et al*, 2009). This is similar to the conclusions in the polyploid persimmon in which the DNA methylation in regulatory sequence controls the production of male and female flowers in genetically male trees (Akagi *et al*, 2016, 2014). Thus, our analysis enabled us to show how multiple gene regulatory pathways are involved in flower development and in meristem maintenance of *S. latifolia*.

### Hermaphrodite flower development affects male fertility and placenta organization

Hermaphrodite development was examined using a combination of WFAD and ISEDS morphology-sensitive detectors, which allowed us to visualize detailed morphological changes at the cellular level (Fig. 1, SI Appendix, Fig. S4). Using these detectors, we obtained unique images of very large samples with high depth of field as well as specific details with high resolution and precision (Neděla *et al*, 2015, 2016), compared to fixation-based electron micrographs of *S*. flowers (Kawamoto *et al*, 2019; Kazama *et al*, 2016; Uchida *et al*, 2003; Koizumi *et al*, 2007). Previous studies examining the infection of *Microbotryum lychnidis-dioicae* in male flowers using conventional scanning electron microscopy showed changes in infected females in earlier flower stages. This revealed the differences between males and females in the onset of teliospore formation (Uchida *et al*, 2003). In addition, infection promoted stamen development in female flowers and elongation of filaments in asexual mutants, as in NT males. Yet, the gynoecium was suppressed within the same asexual flower (Kawamoto *et al*, 2019). In this study, we observed slightly different changes in male and female development. WT male flowers contained a suppressed gynoecium and fully developed anthers with viable pollens (Fig. 1a, SI Appendix, Fig. S5a, i). Hermaphrodite flowers contained a gynoecium with a fully developed ovary and one to three stigmata, suppressed stigma, and a significantly higher number (*P* < 0.001) of aborted pollen grains (Fig. 1d, h, SI Appendix, Fig. S5b–f,i). Nevertheless, the gynoecium in hermaphrodites possessed fewer ovules, which can be influenced by the age of the plant (Yuan & Kessler, 2019). This was avoided by scoring only the first 10 flowers after inflorescence initiation (SI Appendix, Fig. S6, S7).

### Transcriptome profiles and expression changes in hermaphrodites are stable between generations

Rapid transcriptional changes in hermaphrodite flowers inherited through the male germline in *S. latifolia* affected 103 sex-linked and 1,706 non-sex-linked genes in HF1 and 153 sex-linked and 2,468 non-sex-linked genes in HF2 (Table S5). This is similar to the values based on the GO enrichment analysis, in which GO terms related to chemical treatment response reached from 2.44% to 4.49% of downregulated and 2.34% to 4.28% of upregulated genes in hermaphrodites (downregulated in males; Table S9). Thus, our results indicate that expression changes with stable ratio profiles may affect only a small percentage of regulatory genes in a heritable fashion and that only a small subset of genes are up- or downregulated in response to chemical treatment, based on the GO enrichment analysis. In *Arabidopsis*, application of histone methylase and histone deacetylation inhibitors had non-redundant effects that affected only 3–4% of genes (from a total of 7,800 monitored genes), similar to this study (Chang & Pikaard, 2005). Further, the number of affected genes in this study is not different from the number of genes affected by smut fungus (4.7% of non-sex-linked and 7.2% of sex-linked genes up- or downregulated) (Zemp *et al*, 2015).

It is tempting to speculate that a small number of affected genes are inherited by subsequent generations and regulated by an epigenetic mechanism (Janoušek *et al*, 1996; Baubec *et al*, 2014; Niederhuth & Schmitz, 2014). Although ZEB affects global DNA methylation in a dose-dependent and transient manner (SI Appendix, Supplemental Note S6), independent of cytosine context (Nowicka *et al*, 2019; Baubec *et al*, 2009; Griffin *et al*, 2016), epigenetic inheritance from parents with divergent epigenomes permits long-lasting epiallelic interactions (Reinders *et al*, 2009; Zhao *et al*, 2011; Madlung *et al*, 2002). The similar expression pattern between first- and second-generation hermaphrodites shows stable inheritance of the expression profiles for studied genes. The fact that we identified only a small number of histone modification- and DNA methylation-related genes (namely, *DDM1b*, *SUVH4b*, *SUVR4*, *JMJ13*, *JMJ26*, *HAC1c*, *HAF1*, *HDA8*, *HDA9*, *SRT1*, and *SRT2*; Fig. 2) with strong expression differences between males, females, and hermaphrodites supports our conclusion that regulatory pathways related to flower development are key drivers of sex determination in *S. latifolia*. These key drivers will be possibly regulated by the affected histone modification- and DNA methylation-related candidates (She & Baroux, 2014; Vanyushin & Ashapkin, 2011). Nevertheless, the stable expression profiles for most of these genes show that no further changes at the level of epigenetics regulation occur in both observed generations compared to NT males (Fig. 2). Further, the expression of DNA damage and stress-related genes remains the same within the samples (Fig. 2d), as previously shown in *Arabidopsis* (Liu *et al*, 2015; Nowicka *et al*, 2019). The different expression of ABI3 and ERECTA is not surprising, as both genes affect embryo development or inflorescence architecture and must be present if the ovary is formed (Fig. 2f).

The difference in TE expression between males and females is in agreement with previous results showing specific proliferation of those repeats in separate sexes (Puterova *et al*, 2018; Hobza *et al*, 2015). The differences in expression of *Retand*, *Athila*, and *Copia* elements are surprising because TEs are kept inactive in plants by transcriptional gene silencing (TGS), which provides a checkpoint for correct epigenetic inheritance during the transition from the vegetative to reproductive stage, as shown in *Arabidopsis* (Baubec *et al*, 2014; Pecinka & Liu, 2014). Additionally, it was shown that ZEB induces transcriptional reactivation for TEs only in embryonic tissue (before flowering) (Baubec *et al*, 2014). Thus, this result suggests not only that a subset of genes de novo upregulated in *S. latifolia* hermaphrodites is skewing the epigenetic inactivation but also that *Retand*, *Athila*, and *Copia* elements are deregulated with stable epigenetic inheritance, escaping the TGS regulatory pathway (SI Appendix, Fig. S10, Table S8).

### Gynoecium development includes upstream regulator genes

We further classified all GO terms in BP to subclasses based on their specific function (Table S10). From these genes, 4.52% of GO terms (213) belonged to flower development. Because 7,540 GO terms have no classification (Table S9), it is possible that some sequences in the GO categories (and mainly in flower development) are missing. GO annotations are based mainly on references from *Arabidopsis*, a species that shared a common ancestor with *S. latifolia* 180 mya (Kumar & Hedges, 2011). Thus, the predictions from GO ontology may be inaccurate for orthologous sequences, although GO is well suited to cross-species analysis (Primmer *et al*, 2013). To specify the expression profiling of hermaphrodite-related changes, we differentiated five groups consisting of nine clusters (Cl1-9) based on gene expression levels, (Fig. S13). The comparison of sex-linked genes, autosomal genes, and GO terms upregulated in hermaphrodites and GO terms enriched in flowering and GO terms upregulated in HF1 and HF2 yielded 45 sex-linked and 333 autosomal genes related to flower development (Fig. 3a; Table S11). The final comparison of these flower development-related genes with the GO enriched terms (Table S10) and, Group 1 and Group 2 genes (Table S13, SI Appendix, Fig. S13) further specified eight candidate genes, three sex-linked and five autosomal, namely, *HAN*, *ANT*, *GAOX1*, *SPL4, FRL4a*, Z*WIP2*, *YAB1*, and *ATH1* (Fig. 3b). The largest expression differences among profiles were found directly between these genes and their downstream regulated targets (Fig. 4a,b; SI Appendix Fig. S14). Among these candidates, none were previously reported in *S. latifolia* (Kazama *et al*, 2009). All eight candidates determine flower size and morphology (Alvarez-Buylla *et al*, 2010), and the strongest candidate *HAN* has an irreplaceable role in the meristem-to-organ transition, flower organization, and meristem boundary discrimination as found in *Arabidopsis* and other plant models (Hobza *et al*, 2018; Yu & Huang, 2016; Cucinotta *et al*, 2020; Ding *et al*, 2015; Behringer & Schwechheimer, 2015). Thus, we suggest that the *HAN* homolog may directly determine meristem boundaries in developing *S*. *latifolia* flowers based on gene-specific upregulation in hermaphrodites compared to NT male flowers. This is supported also by previous evidence examining the expression of *S. latifolia* floral MADS-box genes in early developmental stages using *in situ* hybridization. The authors suggested that gynoecium-suppressing genes may act on the factors that determine the boundaries of the whorls (Hardenack *et al*, 1994). If this is true, *HAN* is an ideal candidate, either promoting or suppressing gynoecium development through regulation of *YAB1*, *JAGGED*, and *CKX3* and its own autoregulation (Yu & Huang, 2016; Ding *et al*, 2015).

Among the candidates, the *JAGGED* transcription factor gene is directly regulated by HAN and plays an important function in stamen and carpel shape. *JAGGED* together with *AS1 transcription factor* and *AS2 protein* (*Asymmetric leaves*) functions in cell proliferation, repressing the boundary-specifying genes in sepal and petal primordia in *Arabidopsis* (Xu *et al*, 2008; Dinneny *et al*, 2006). Notably, *HAN* also promotes the expression of *Cytokinin dehydrogenase 3* and indirectly represses the expression of *KNOX1* genes through *YAB1*, thus promoting cell differentiation through repression of gibberellin inhibitors (Kumaran *et al*, 2002; Yu & Huang, 2016). ANT directly affects the expression of *REVOLUTA* (*REV1*) and *PERIANTHIA* (*PAN*), which determine the floral organ number and patterning (Otsuga *et al*, 2001; Das *et al*, 2009). WUSCHEL (WUS), LEAFY (LFY) and PAN among others, positively regulate *KNUCKLESS* which in turn repress expression of *WUS* to terminate the stem cell niche once a limited number of organs have been achieved (Alvarez-Buylla *et al*, 2010). In addition, PAN regulates *AGAMOUS*, a C class MADS-box gene (Das *et al*, 2009; Alvarez-Buylla *et al*, 2010). *SUPERMAN*-*like* (*SUP*), previously reported as a negative regulator of male organ development (Kazama *et al*, 2009; Fujita *et al*, 2019), was also specifically upregulated in hermaphrodites compared to males. In *Arabidopsis*, *SUP* is negatively regulated by YAB1 (Yu & Huang, 2016; Sawa *et al*, 1999). Another MADS-box protein AGL42, regulated by SPL4, controls the flowering time and acts through a gibberellin-dependent pathway (Dorca-Fornell *et al*, 2011). Unsurprisingly, the transcription factor IND, regulated by ZWIP2, was only upregulated in hermaphrodites and expressed in females. This transcription factor specifies the development of the valve margin and plays a role in the cell type differentiation in fruit dehiscence in *Arabidopsis* (Liljegren *et al*, 2004). Among the MADS-box genes involved in flower development, homologs of *PISTILLATA* (*SLM2*) and *APETALA3* (*SLM3*), genes specifying stamen development, were expressed at lower levels in female flowers compared to males, or HF1 and HF2. Because *AGAMOUS* is also important during stamen and anther development, its high expression is not surprising. Nevertheless, D class MADS-box gene homologs (*STKa*, *STKb*, *STKc*, and *STKd*) were strongly expressed in hermaphrodite and female flowers compared to males.

Previous experiments examined the role of multiple genes, including ABC MADS-box genes, in flower development and organization in *S. latifolia* (Hobza *et al*, 2018). The expression of MADS-box flowering genes in this work differed between males and females in B- and D-class MADS-box genes and for D-class MADS-box genes in hermaphrodites compared to males (Fig. 4b, SI Appendix, Fig. S14). Nevertheless, there is an elementary difference between dioecious plants in which the organs of the opposite sex are completely suppressed and those possessing rudimentary organs (Cronk & Müller, 2020). MADS-box genes may be considered candidates only if the organs of the opposite sex are entirely missing (Cronk & Müller, 2020). The situation is opposite in *S. latifolia*, which still possesses a rudimentary gynoecium in males (Fig. 1a) and suppressed rudimentary stamens in females (Fig. S4a). Therefore, the presence of a vestigial gynoecium suggests that upstream regulatory genes have an important role in carpel development (Cronk & Müller, 2020; Graham *et al*, 2003). This possibility is supported by the differential expression of *HAN*, *ANT*, *GAOX1*, *SPL4*, *FRL4a*, Z*WIP2*, *YAB1*, and *ATH1* in *S. latifolia* hermaphrodites (Fig. 3b, Fig. 4a). We hypothesize that sex determination takes place after the flower meristem is partioned into four concentric whorls of primordial cells, and that these key regulators are among the first genes to be affected by the sex determination factor, which is yet to be identified.

Furthermore, it remains unclear how epigenetic modifications regulate sex and flower morphology in *S. latifolia.* Further studies should focus on the regulatory mechanism of their expression. The morphological changes after application of chemical genetics show that sex determination in *S. latifolia* is influenced by an epigenetic-related pathway, as previously reported in *Diospyros kaki* (Akagi *et al*, 2016). It is also tempting to speculate that DEGs related to histone and DNA methylation and histone acetylation may play a role in sex determination mechanisms. This should be further tested in relation to the specific function of the sex-linked candidates *HAN*, *ANT*, and *GAOX1*. Finally, it is possible to use genetic transformation and targeted mutagenesis protocols that are now available for *S. latifolia* (Hudzieczek *et al*, 2019). Such approaches simplify further studies of each of the candidate genes and their functions in relation to sex determination. The identification of female promoting genes in this study provides insight into the complexity of flower development, contributing to our understanding of the *S. latifolia* sex determination mechanism and female flower development.

## Acknowledgements

This project was supported by the Czech Science Foundation (no. 19-02476Y and no. 19-03909S) and by the Czech Academy of Sciences [RVO:68081731]. We thank the EMBL Genomics Core Facility (EMBL, Heidelberg) for transcriptome sequencing. Computational resources were supplied by the project "e-Infrastruktura CZ" (e-INFRA LM2018140) provided within the program Projects of Large Research, Development and Innovations Infrastructures. We would like to thank Plant Editors for the English-language correction.

## Author information

These authors contributed equally: Vaclav Bacovsky, Radim Cegan

## Contributions

V.B., R.C., and R.H. conceived the study and experimental design. V.B. did all the experimental work (and associated statistics) and prepared the plant material for sequencing and samples for FE-ESEM. R.C. did the data analysis and ran all bioinformatics tools, analyzed the data, and prepared all the bioinformatics tables and figures with input from the other authors. R.H., L.S. and V.H., analyzed the floral development genes with input from the other authors. E.T. and V.N. did all the FE-ESEM observations and figure post-processing. V.B. led the effort to prepare the library and produce the the sequencing data. V.B., R.C., and R.H. wrote the main text of the manuscript with input from the other authors.

## Ethics declarations Competing interests

The authors declare no competing interests.

## Data deposition

The transcriptome sequencing data are deposited in the European Nucleotide Archive (https://www.ebi.ac.uk/ena/browser/home) under accession PRJEB36078.

